# Tree Growth, Contraction, and Recovery: Disentangling Soil and Atmospheric Drought Effects

**DOI:** 10.1101/2025.04.24.650406

**Authors:** Erez Feuer, Yakir Preisler, Eyal Rotenberg, Dan Yakir, Yair Mau

**Affiliations:** The Institute of Environmental Sciences, The Hebrew University of Jerusalem, Rehovot 76100 Israel; Institute of Plant Science - Agricultural Research Organization - Volcani Institute Rishon LeZion 7505101, Israel; Department of Organismic and Evolutionary Biology, Harvard University, Cambridge, MA, USA; Earth and Planetary Science Department, Weizmann Institute of Science, Rehovot, Israel

**Keywords:** recovery, resistance, dendrometer, dry forest, growth rate, compound drought

## Abstract

We investigate how soil and atmospheric droughts jointly impact tree growth and recovery dynamics in a semi-arid pine forest, leveraging high-resolution stem diameter variation data and an irrigation experiment. The irrigated plot, where soil drought was mitigated, served as a benchmark to isolate the effects of atmospheric drought and distinguish them from the compound drought conditions experienced by control trees. Using a suite of tools based only on stem diameter variation, we identified growth modes that vary in accordance with soil water availability. Control trees showed negligible growth during the dry season but rapidly recovered with the onset of the wet season, matching the baseline growth rates of the irrigated trees, suggesting minimal compromise in hydraulic functioning. Our main finding is that heatwaves consistently depress stem-expansion rates, regardless of treatment. However, during the dry season, this negative impact diverges sharply between the treatments. Because irrigated trees benefit from a hydraulic buffer supplied by ample soil water and thus retain a positive growth baseline, the depression merely slows their expansion, whereas control trees already near zero are driven into net contraction. These findings offer new understanding of how trees balance growth, contraction, and recovery under varying drought conditions, revealing the pivotal role of soil water in shaping drought responses across seasons. As climate change intensifies the frequency and severity of drought events, this knowledge is critical for anticipating shifts in tree growth and resilience.

## 1 Introduction

Large tree-mortality events have been recorded across the globe for many decades (Senf et al., 2018; Yu et al., 2019; Powers et al., 2020; Hammond et al., 2022). Atmospheric and soil droughts are two major stressors impacting tree growth and survival, often linked to shifts in rainfall patterns and intensifying atmospheric dryness (Breshears et al., 2013; Trenberth et al., 2014; McDowell et al., 2022). Atmospheric drought relates to periods when the air is hot and dry, and it is usually quantified by high vapor pressure deficit (VPD) values. Trees’ response to high VPD values are often associated with decreased tree carbon reserves, decreased growth, and higher risk of hydraulic failure, among others (Novick et al., 2024). Soil drought denotes periods of time of low soil water content, usually brought about by low precipitation. In the coming decades, we expect worsening of these two kinds of droughts, regarding their intensity, duration and frequency characteristics, posing an even greater threat to tree survival (Meehl and Tebaldi, 2004; Dai, 2013; Perkins-Kirkpatrick and Gibson, 2017; Chiang et al., 2021; Dore, 2005; Xu et al., 2019). Still worse, the compounding of soil and atmospheric droughts—when both occur together—creates a more complex and intensified stress regime, highlighting the need for a deeper understanding of their combined impacts on ecosystems (Zhou et al., 2019; Yin et al., 2023; Shekhar et al., 2024).

As drought conditions intensify, trees increasingly rely on internal water reserves to sustain essential physiological functions. Drawing on these reserves enables trees to maintain critical processes like photosynthesis, transpiration, and osmoregulation while preventing hydraulic failure and loss of turgor (Meinzer et al., 2009; Steppe et al., 2015; Preisler et al., 2022). As soils dry and canopy transpiration outpaces root water uptake, trees draw on internal water reserves, which gradually deplete. This decline in water status restricts growth, reduces carbon sequestration, and increases vulnerability, eventually raising the risk of mortality (Steppe et al., 2015; Peters et al., 2021; McDowell et al., 2022). Thus, water storage stands out as a crucial component, supporting tree functioning in the face of escalating drought pressures.

The dynamics of stem growth and internal water storage have been increasingly studied through continuous, high-resolution stem diameter measurements using dendrometers. Dendrometers offer the advantage of real-time monitoring, providing insights into both water status and growth (Zweifel et al., 2016). Dentrometry-based metrics can reveal how water storage supports immediate tree functioning while also illustrating the long-term depletion of water reserves (De Swaef et al., 2015). For instance, analysis of stem diameter variations were used to derive early warning indicators of tree death (Preisler et al., 2020; Andriantelomanana et al., 2024). Zweifel et al. (2016) proposed two metrics: irreversible stem expansion (GRO), which quantifies the historical maximum stem diameter and indicates growth through wood formation, and tree water deficit (TWD), defined as the difference between GRO and the actual stem diameter, which reveals temporary stem shrinkage as trees tap into stored water reserves. This framework has provided useful concepts to talk about tree growth (Zweifel and Häsler, 2001), water potential (Dietrich et al., 2018), stomatal conductance (Ziegler et al., 2024), water use strategies (Sánchez-Costa et al., 2015) and the effects of elevated VPD on stem shrinkage (Salomόn et al., 2022).

However, a significant question remains largely unanswered: how to disentangle the influence of atmospheric and soil drought on plant growth and resilience (Novick et al., 2024). These two types of drought often occur together, compounding their effects (Yin et al., 2023). Consequently, the same negative effects on trees associated with high VPD are also associated with low soil water content, making it difficult to determine the individual contribution of each factor to the overall stress on tree growth. Zweifel et al. (2005) proposed a phenomenological model to deal with this question, showing that physiological responses to VPD depend on soil water content. Preisler et al. (2023) found that when eliminating soil drought, even under extreme VPD conditions (> 5 kPa), Aleppo pine trees will maintain high assimilation despite the extreme conditions. Additionally, a recent study by Vargas Zeppetello et al. (2023) suggests that VPD has been overstated in its impact over ecosystem evapotranspiration, and that soil moisture is the most relevant state variable to understand surface water conductance. These findings underscore the need to reevaluate the relative roles of atmospheric and soil drought, with growing evidence pointing to soil moisture as a key driver of plant resilience and ecosystem function.

Our goal here is to reveal how tree growth is impacted by soil and atmospheric droughts. We analyzed stem diameter variations from an irrigation experiment in a semi-arid pine forest—an expansion of the experiment covered by Preisler et al. (2023). We developed new methods for heatwave characterization and introduced a suite of tools for assessing growth and resilience. These methods aim to deepen our understanding of tree functioning under complex drought conditions and to provide insights into the fate of forested ecosystems in a warming and more variable climate.

## 2 Methods

### 2.1 Study site and experimental design

The study was conducted at the Yatir forest, a semi-arid forest located at the northern edge of the Negev desert in Israel (31.345°N, 35.052°E, 550–700 m elevation). The managed forest, predominated by Allepo pine (*Pinus halepensis*), was planted in the 1960s. The climate in this region is Mediterranean, characterized by hot and dry summers (mean July temperature: 25°C) and mild, wet winters (mean January temperature: 10°C) with mean-annual rainfall 229 mm (calculated for the years 2000–2022), concentrated in months Dec–May. The forest is subjected to several dry and short heatwaves throughout the year, where VPD can reach values of up to 6.5 kPa (Tatarinov et al., 2016).

Two adjacent plots were established: a control plot with natural soil moisture levels and a plot that received additional irrigation. These plots, which are separated by a 30-meter-wide buffer zone, have similar topographic and ecological features, including slope, tree density, and tree age. Out of the 20–30 trees in each plot, seven in the control plot and six in the irrigation plot were equipped with band dendrometers. Irrigation began on May 14th, 2017, with water supplied daily via drip irrigation according to the monthly average potential evapotranspiration (PET) data from the Israel Meteorological Service. In 2020 and 2021, the irrigation rate was reduced to 80% and 50% of PET, respectively. Additional details on the experimental site at the Yatir forest can be found in previous publications (Grünzweig et al., 2007; Tatarinov et al., 2016; Preisler et al., 2019, 2023). Unlike the agricultural context, where irrigation is primarily used to boost yields, the aim of this setup was twofold. First, it allowed us to distinguish between the effects of compound droughts in the control plot and those of atmospheric droughts alone in the irrigated plot, where soil drought was mitigated. Second, the irrigated plot served as a soil drought-free benchmark against which the growth of the control trees could be compared.

### 2.2 Season classification

The precipitation regime in the Yatir forest imposes a clear seasonal pattern in soil moisture throughout the year (Preisler et al., 2019). In order to capture qualitative differences of tree behavior as a function of soil drought, we divided the year into four seasons: wetting, wet, drying and dry. The wetting season starts on the first rainfall event of the hydrological year, and ends when precipitation reaches 35 mm, which is approximately the rain needed to pass the transpirable soil water content threshold for the Yatir forest (Klein et al., 2014; Preisler et al., 2019). The wet season follows, lasting until the date corresponding to 80% of that year’s total precipitation. We found that 80% is approximately the time of divergence of growth rate between the trees in the control and irrigated plots, as described in the Results section. The drying season then begins, lasting for 60 days. Finally, the dry season follows, lasting until next year’s wetting season.

### 2.3 Heatwave classification and characterization

In this study we use dendrometer data to assess whole tree water status. Water status is the outcome of tree water balance, whose input is soil water uptake and output is transpiration. Transpiration’s major driver are elevated atmospheric demand events, characterized by high VPD values, hereafter simply “heatwaves”. When discussing heatwaves, we have two main goals: identifying their duration, and attributing to each of them a severity score (see Rez et al. (2024) for an extended discussion).

As will be shown in the Results section, there are rich seasonal dynamics to the tree water status. In order to fully capture these dynamics, we chose to classify season-specific heatwaves by analyzing instantaneous VPD percentiles rather than absolute values (Robinson, 2001). We calculated VPD percentiles using 26 years of available 10-min frequency weather data for the nearby meteorologic station (Shani station; Israel Meteorological Service). The calculation was done by pooling (aggregating) all VPD values from a two-week window around one-hour intervals (Fischer and Schär, 2010). For example, if the VPD percentile at 10:30 on April 14th, 2019 is 80, it means that the VPD value at that time is in the 80th percentile of all VPD values from 10:00–11:00 in the April 7–21 span in the years 1997–2023. This method allows us to capture the shifts in VPD relative to the distribution with sub-daily resolution. Time periods that exhibited VPD percentile values above 50, allowing for up to a 6-hour tolerance in dips below 50, are considered potential “heatwave events”. This pooling of heatwaves separated by short intervals is called inter-event time method, and it is standard in drought-identification practices in hydrology (Tallaksen et al., 1997). We are left with a large number of identified potential events, with interesting edge cases. There will be short events that last for just a few hours but are intense (high VPD percentiles), and other much longer events of many days, whose VPD percentile values hover just above 50. In order to prioritize VPD percentile strength over long duration, we computed each event’s severity as

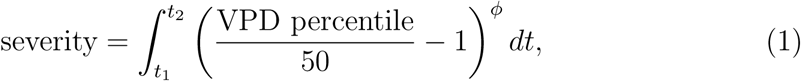

where time, and therefore severity, are in days. The time instants t_1_ and t_2_ denote the beginning and end of a given event. A full day at VPD percentile = 50 will yield a severity of 0, while a full day at VPD percentile = 100 will yield a severity of 1. The parameter ϕ controls how much more high-VPD percentile values count with respect to lower values. For ϕ = 1, VPD percentile = 100 counts twice as much as 75, while for ϕ = 2 it counts four times as much. In this paper we chose ϕ = 2.5, and potential heatwave events will be considered as such when their severity surpass the threshold of 0.03. To be sure, there will always be some arbitrariness in defining a heatwave event, but we tried to strike a balance between complexity and generality. In order to be more strict or lax regarding one’s definition of a heatwave event, it suffices to choose other values for ϕ and the severity threshold. See Fig. S1 for a representation of heatwaves and their severity during an eight-month interval. Normalized severity is defined as the severity divided by the duration of the event (i.e., divided by t_2_ *−* t_1_), therefore it is also interpreted as the mean daily heatwave severity.

### 2.4 Flux measurements

Flux measurements were performed using the eddy covariance technique. An instrumented tower operates at the center of Yatir Forest since 2000, following Euroflux methodology (Aubinet et al., 1999; Grünzweig et al., 2003). The system uses a 3D sonic anemometer (Gill R-50; Gill Instruments, Lymington, UK) and a closed path infrared gas analyser (LI-7000, LI-COR, Lincoln, NE, USA) to measure H_2_O and CO_2_ fluxes. The tower is also equipped with a full meteorological station, providing measurements of air temperature, relative humidity, precipitation and wind speed. Raw data were analyzed with the use of EddyPro software (Li-Cor, Lincoln, NE, USA) in half-hour time step. Outliers and spikes were detected and removed with the use of the double-differenced time series, using the median of absolute deviation about the median (Papale et al., 2006). Gap filling was performed with the marginal distribution sampling (MDS) methodology and flux partitioning was based on the night-time fluxes method (Reichstein et al., 2005). The above post-processing analyses were performed with the use of the REddyProc R-package (Wutzler et al., 2018).

### 2.5 Dendrometer measurements

Each tree in this study was equipped with a band dendrometer (EMS, Brno, Czech Republic) at breast height (1.3 m) measuring variations in stem circumference. These measurements were divided by π to obtain variations in diameter at breast height (DBH). In this study, we aimed to identify shifts in DBH dynamics driven by atmospheric events, beyond the usual daily expansion and contraction of the trunk. To isolate this broader trend, we applied a 1-day rolling average, effectively filtering out the daily signal while retaining the underlying temporal pattern. From DBH alone we derive a number of useful metrics: absolute growth (GRO), absolute growth rate (GRO rate), tree water deficit (TWD), resistance ratio (Ω), tree recovery index (T-REX), and DBH slope, to be detailed below.

#### GRO

The absolute growth is the historic maximum DBH, called GRO in the zero growth model proposed by Zweifel et al. (2016).

#### Growth rate

Growth rate is a non-negative quantity, defined as the smoothed time derivative of the GRO, obtained by a 45-day Savitzky-Golay differentiation filtering of order one. Because GRO consists of plateau periods interrupted by sudden jumps, smoothing is necessary to minimize sharp spikes in the derivatives, resulting in a more stable, easy to interpret time series. It is important to note that growth rate relates solely to the derivative of GRO, that is, it considers only the dynamics of the tree stem. To be sure, different parts of the tree, such as needles and stem, have their own seasonal growth patterns (Maseyk, 2006; Maseyk et al., 2008).

#### TWD

Tree water deficit (TWD) was calculated as the difference between the GRO and the DBH (Zweifel et al., 2016).

#### DBH Slope

The DBH slope represents the average rate of change in stem diameter over a given time window, serving as an indicator of expansion or shrinkage. It is calculated as the slope of a least-squares linear fit.

#### Ω

The resistance ratio (Ω) is the proportion of days (unitless), within a given time window, that registered tree diameter expansion. As such, Ω ranges between 0 and 1. High Ω values signify a tree’s ability to withstand disturbances over that period, while low values indicate susceptibility. We calculated Ω for all trees using a rolling window of 45 days.

#### T-REX

Tree REecovery indeX (T-REX) evaluates tree recovery abilities from stem contractions. T-REX is calculated thus. First, the GRO and TWD are downsampled to daily frequency using the first value of each day. T-REX is zero whenever the DBH coincides with the GRO (i.e., TWD is zero). It is incremented by 1 for each day in which the TWD increases, therefore the units of T-REX are day. If TWD decreases, T-REX takes the value it had when TWD last reached this level. A step-by-step demonstration of the calculation using a synthetic DBH time series is illustrated in Fig. 1. T-REX values represent the effective number of days a tree has experienced stem shrinkage since it last registered absolute growth. An increasing T-REX indicates sustained contraction and a lack of recovery, whereas a rapid decline reflects a return to growth and successful recovery. As such, T-REX serves as a recovery index: higher values denote lower recovery capacity. Under similar environmental stress, trees with higher T-REX values are those that recovered less along the way. We propose a recursive algorithm for computing the T-REX in the Supporting Information, this algorithm was used for all T-REX calculation throughout this study.

**Figure 1:**
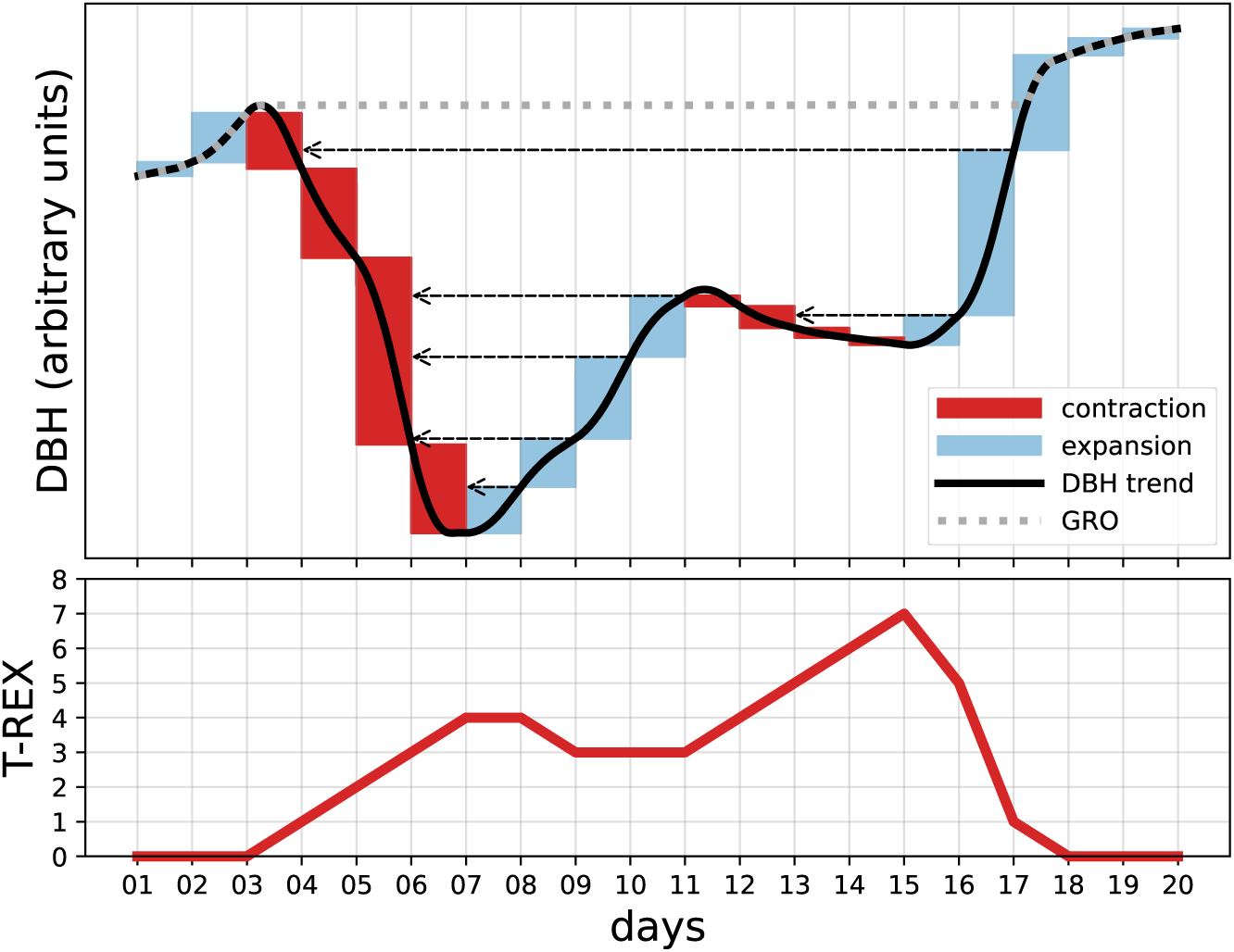
Calculation of Tree REcovery indeX (T-REX). T-REX captures daily trunk recovery ability, rising by 1 for every day with increased shrinkage and resetting when expanding to a prior state. **Walk-through**: The dotted gray line shows the GRO curve (Zweifel et al., 2016), which can be understood as the historical maximum diameter. Whenever tree diameter (black line) coincides with GRO, T-REX equals zero. Whenever the stem contracts over a span of one day (red blocks), T-REX is incremented by one (thus T-REX’s unit is ‘day’). Upon stem expansion (blue blocks), T-REX either stays the same or goes down in value. Its updated value is the same as the T-REX for the previous day with equal diameter at breast height (DBH) (see dashed gray arrows).

### 2.6 Growth Modes

We classified tree growth into six modes, shown in Fig. 2. The modes are determined for a given time window by their Ω and T-REX values. We differentiate between two main cases: modes with an increasing GRO (left half of the figure) and those with a stagnant GRO (right half). Among the increasing-GRO modes, as Ω decreases there is a gradual shift from full growth, to disrupted growth, and then to spurt growth. We use the term full growth to describe stable and uninterrupted growth, while disrupted growth reflects active growth with transient setbacks, and spurt growth captures brief bursts of growth following prolonged periods of zero growth. Conversely, in stagnant-GRO modes, changes in the T-REX slope—from positive, to near-zero, to negative—reflect a transition from contraction to standstill, and finally to recovery. We refer to contraction as a phase of ongoing shrinkage, standstill as a state where contraction and expansion alternate without net growth—reflecting a balance between water loss and partial replenishment—and recovery as a period of expansion that restores previously lost volume, without yet reaching the point of renewed growth.

**Figure 2:**
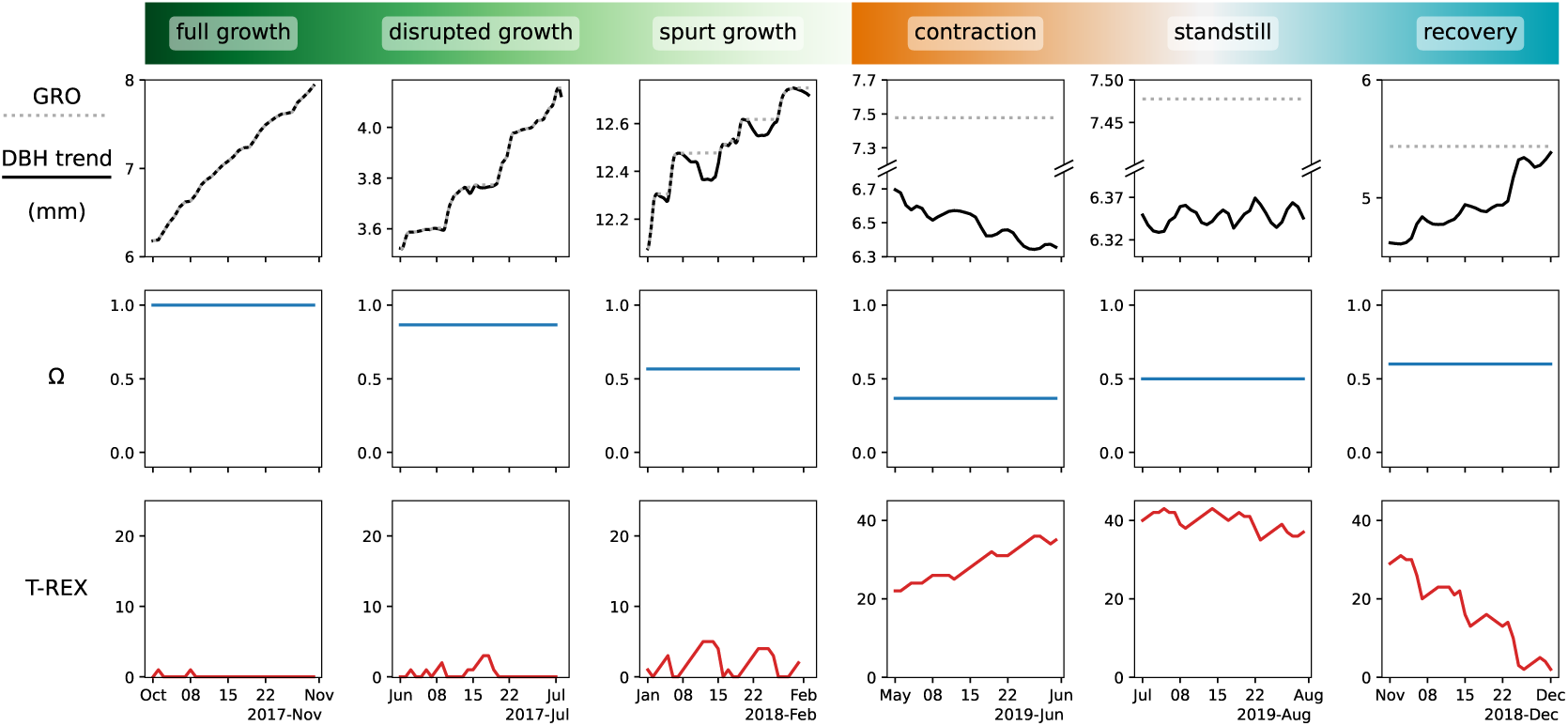
Growth mode classification. Each column illustrates an example of a growth mode, based on one month of real data. Rows show the DBH trend (solid black) and GRO (dotted gray), as well as the corresponding Ω (blue) and T-REX (red) values for that one-month window.

### 2.7 Atmospheric events and response

The complete climatic time series was segmented and categorized into four types of atmospheric events: heatwave, post heatwave, rain, and post rain. We have already discussed heatwave classification above. Rain events begin when precipitation is first recorded and end when it stops. Rain events occurring on the same day or on contiguous days were merged into a single rain event, starting with the first precipitation and ending with the last in that combined period. All periods following a heatwave or rain event that were not otherwise classified were labeled as post heatwave and post rain, respectively. We quantified the tree response to these short atmospheric events using the DBH slope.

### 2.8 Statistical analyses

#### Linear mixed-effects model

The connection between heatwave severity and DBH slope was determined using a linear mixed-effects model. In this model, heatwave normalized severity, treatment (control or irrigated), and season were chosen as fixed effects, and individual tree IDs were included as a random effect. We examined both the two-way and three-way interactions involving heatwave normalized severity. We focused on only two seasons—the dry and the wet—as they represent the extremes in soil moisture conditions and exhibit less variability within each season. We chose the wet season as the default season because, during that time, water is generally available in the soil and similar across both plots. As the name suggests, control was used as the default treatment in this analysis. Fit was done using the statsmodels Python package. Marginal and conditional R^2^ were calculated according to Nakagawa and Schielzeth (2013).

#### Permutation test

Quantifying the difference between the DBH slope of the control and irrigated groups requires special care. The responses of all trees to all atmospheric events are not independent — they consist of the responses of a few unique trees to many unique events, which are identical across all trees.

To compare the responses between trees in the control and irrigated plots, we used the median of the differences in their responses across all atmospheric events. Specifically, for each unique event (e.g., rain or heatwave), we first calculated the median response of the trees in the control plot and the median response of the irrigated trees. We then computed the difference between these two medians for each event. After obtaining the differences for all events, we took the median of those differences to quantify the overall response difference between the two groups. Our null hypothesis was that this median difference would be zero, meaning that the median responses of irrigated and control trees would be the same across all atmospheric events.

Since each atmospheric event affects all trees in both groups, responses for a single event are dependent because they come from the same trees. To account for this dependency, we computed p-values using an exact permutation test. In this test, we shuffled the treatments of the trees across all atmospheric events. This means that each permutation shuffles the treatment of all 13 trees (7 control and 6 irrigated), and this new labeling is applied to all atmospheric events. This approach preserves the dependence structure within each tree and accurately reflects how consistent tree responses affect all comparisons.

The number of valid permutations was the total number of permutations, excluding permutations within the same group, as this would produce identical results. The exact number can be calculated using the binomial coefficient, in this study— n*_c_* = 7 control trees and n*_i_* = 6 irrigated trees, that is 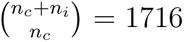. We then adjusted the p-values using the Benjamini–Hochberg procedure to account for the false discovery rate (Benjamini and Hochberg, 1995).

## 3 Results

### 3.1 Growth Rate

Irrigated trees exhibited growth rates strongly linked with average temperatures and sunlight hours: maximum growth rates occurred when temperatures were high and the photoperiod was long, and vice versa (see Fig. 3). As the amount of irrigation decreased, the spread in growth rates among irrigated trees also decreased. In 2021, when irrigation was further reduced from 80% to 50% of PET, growth rates noticeably declined, with the peak mean growth rate being roughly half of that observed in previous years with higher irrigation levels.

**Figure 3:**
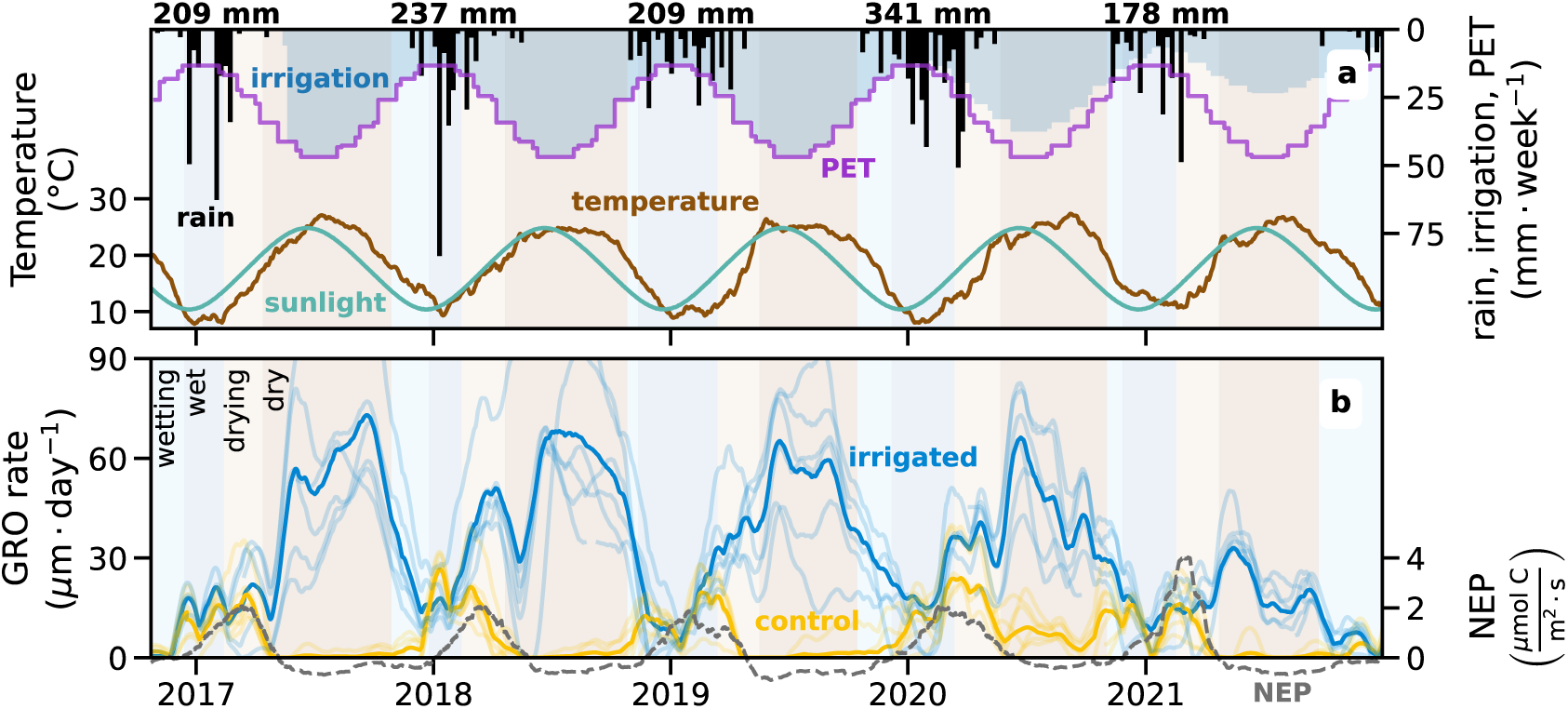
Growth rates of trees in the irrigated and control plots vary seasonally, responding to temperature, photoperiod, and water availability. Irrigated trees are by design not water-limited, peaking with high temperatures and long photoperiods, while control trees show water-limited growth, peaking during the wet season with high water availability. **Panel a:** Climate and irrigation data for the Yatir forest. Black numbers above the panel indicate annual precipitation for each hydrological year. Irrigation treatment started on May 14th, 2017, and was set to the monthly average PET. In 2020 and 2021, irrigation was decreased to 80% and 50% of PET, respectively. The temperature curve represents the 30-day rolling mean of the air temperature at 2 meters. The sunlight curve indicates the number of daylight hours per day, with a minimum of 10 hours and a maximum of 14 hours. **Panel b:** Growth rates of irrigated and control trees, shown as the GRO rate. Curves represent the derivative of the 45-day rolling mean of the GRO, with thin lines indicating individual trees and thick lines representing their mean (control: n*_c_* = 7, irrigated: n*_i_* = 6). NEP data was acquired from the flux tower in Yatir, representing the entire Yatir forest, which is growing under the same conditions as the control trees. Wetting, wet, drying, and dry seasons are determined according to the precipitation regime as described in the Methods section.

Trees in the control plot showed peak growth rates during the wet season with relatively low temperatures and light hours but high water availability. During that time, growth rates were comparable to the mean growth rates of the irrigated trees. The peak in growth rates coincided with the peak in Net Ecosystem Productivity (NEP), as measured by the adjacent flux tower. During the drying and dry periods, as soil water availability decreased, control growth rates declined to near-zero. This decline also corresponded with the NEP trend. The dry season of 2020 was unique, as control growth rates were higher than the previous years. That year also had elevated precipitation levels of 341 mm. Whenever the wetting period started, within a window of 1–2 months, the growth rates of the control trees rose to match those of the irrigated trees, and remained similar until the drying period.

A closer look at the growth differences and similarities between the control and irrigated plots is provided in Fig. 4 using the Ω and T-REX metrics. The resistance ratio Ω of irrigated trees mirrors their growth rate trend, with high values (near 1) during the dry season and lower values (around 0.5) during the wet season. In contrast, the control plot maintained Ω values around 0.5 over the entire five-year period. On a seasonal scale, however, trees in both plots showed coinciding Ω values during the wet season. Notably, in the later years of the experiment (2020 and 2021), when irrigation amounts were reduced, the Ω values of the irrigated plot decreased, approaching those of the control plot.

**Figure 4:**
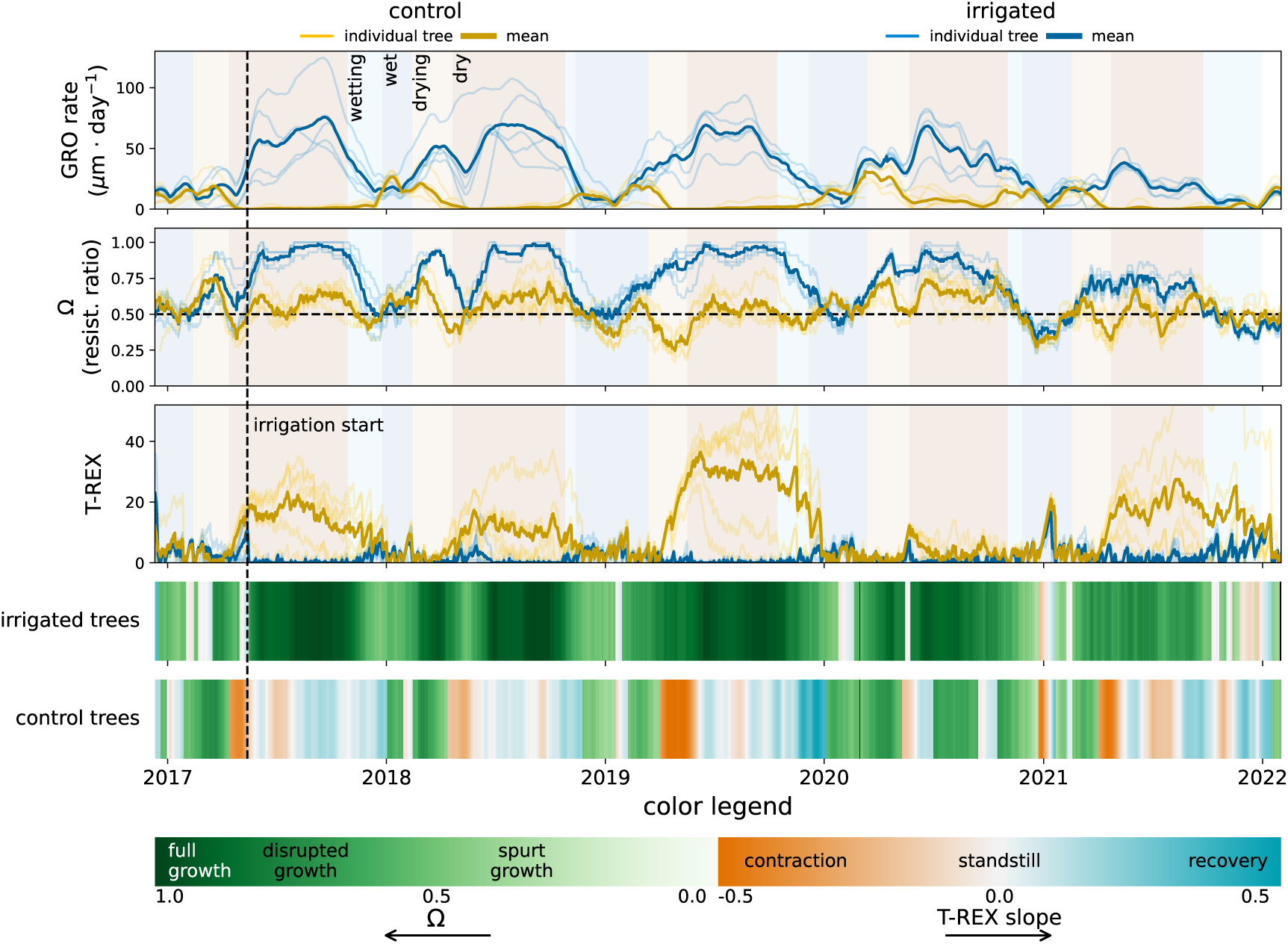
Resistance ratio, T-REX, and growth mode dynamics. The top three panels show the dynamics of GRO rate, Ω, and T-REX. Thin lines indicate individual trees; thick lines represent the mean (control: n*_c_* = 7, irrigated: n*_i_* = 6). GRO rate and Ω were calculated using a 45-day rolling window. The bottom panels show growth mode dynamics for irrigated and control trees. Stagnant-GRO modes were defined as periods when mean T-REX exceeded 7 days and ended once it dropped below 2 days. To better capture the onset, these modes were backdated by 7 days from the threshold crossing. Color shading (orange to blue) reflects the T-REX slope over the 45-day window. All other days were classified as increasing-GRO modes, with shading from white to green representing increasing Ω values.

T-REX showed strong seasonal fluctuations in the control plot, with high values during the dry season (reaching above 30) and low values during the wet season (near 0). The opposite pattern was observed in the irrigated plot, where T-REX peaked during the wet season and was negligible during the dry. Here too, the two treatments showed similar T-REX values during the wet season. The relatively wet hydrological year of 2020 was followed by an anomaly in which T-REX registered low values for control trees during the dry season.

Constructing a time series of growth modes reveals a clear picture of how each plot responded over the course of the experiment (see Fig. 4). Irrigated trees remained in increasing-GRO modes for most of the year, typically showing full growth during the dry season and shifting toward spurt growth in the wet season. As irrigation levels declined, full growth became less frequent. In contrast, control trees exhibited minimal growth throughout the year. They spent most of the time in stagnant-GRO modes—cycling through contraction during the dry season, followed by standstill, and then recovery during the wetting and wet seasons—showing only brief periods of active growth.

### 3.2 Seasonal Responses to Atmospheric Events

Aiming at revealing correlations between growth dynamics and both seasons and atmospheric events, we analyzed the statistics of change in DBH slope during these events, as shown in Fig. 5, and as detailed in Methods. Each panel shows probability distributions of the DBH slope for each treatment (control in yellow, irrigated in blue), for all combinations of seasons and atmospheric events, excluding the rain and post-rain events in the dry season, due to insufficient data for a meaningful statistical analysis. Stem expansion and contraction correspond to positive and negative DBH slope values, respectively.

**Figure 5:**
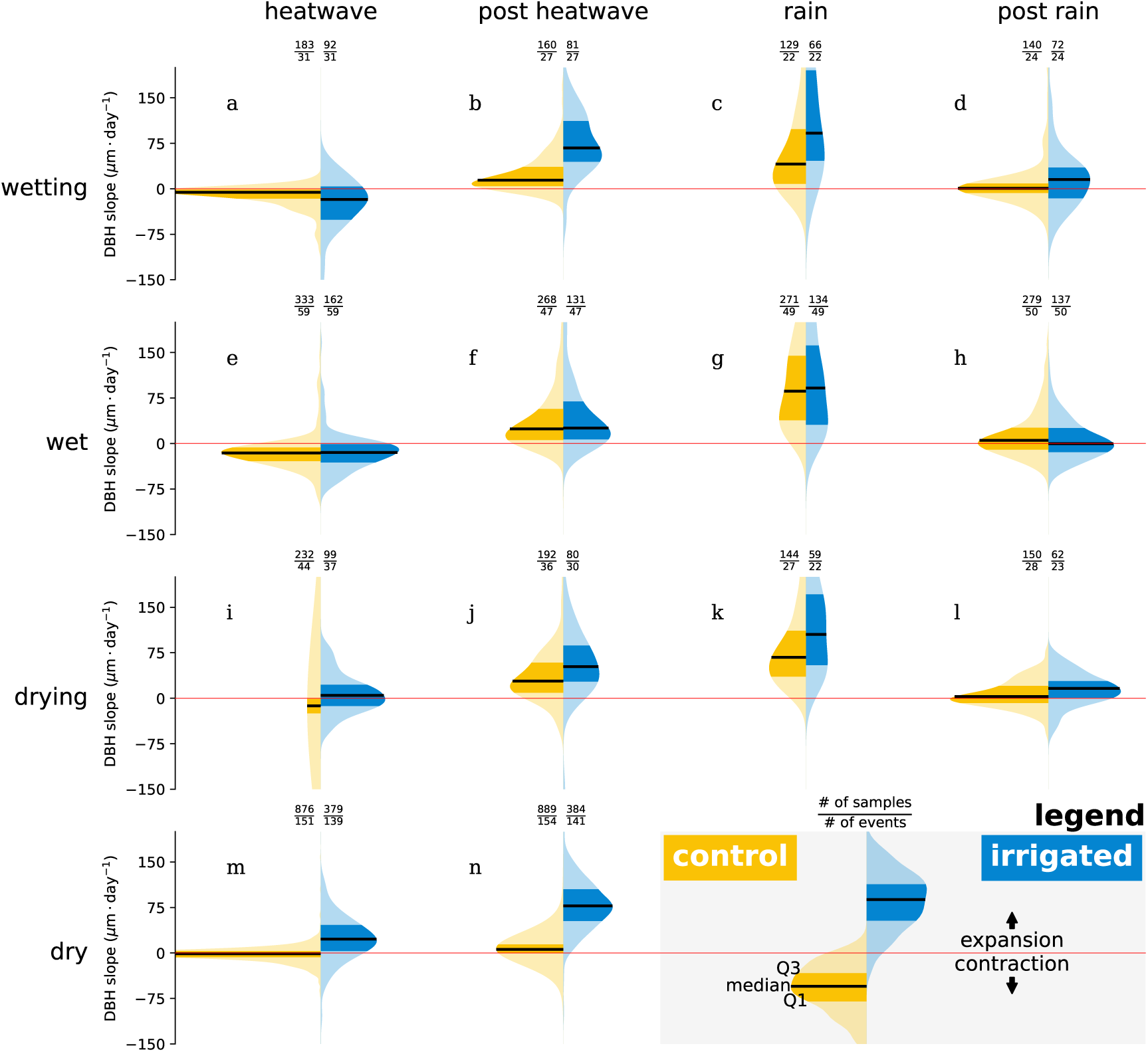
Control and irrigated trees respond indistinguishably during the wet season across all atmospheric conditions. The figure compares the distribution of the DBH slope across all atmospheric events (heatwave, post heatwave, rain, post rain) and seasons (wetting, wet, drying, dry) between control and irrigated trees. As detailed in the legend in the bottom right, probability densities for irrigated trees are colored blue, and control are colored yellow, both plotted in the vertical direction, and arranged in a configuration similar to violin plots. The median values are marked with a black line, while the interquartile range is shaded with a higher color saturation. The fraction of number of samples to number of events is indicated over each distribution.

Across most combinations of seasons and atmospheric conditions, the control and irrigated trees showed statistically distinct DBH slope distributions (see Table S1. However, during the wet season, under all atmospheric conditions, the two treatments were not significantly different, with consistently high p-values indicating similar responses. A similar lack of difference was also observed during post-rain events in the drying season. During rainfall events, both control and irrigated trees exhibited stem expansion across all seasons. The irrigated trees generally showed higher median expansion rates than the control trees. Under heatwave conditions, control trees consistently showed negative median DBH slope values, indicating stem contraction. The irrigated trees under heatwaves, in contrast, mostly contracted during the wetting and wet seasons but expanded during the drying and dry seasons. When comparing heatwave and post-heatwave conditions within each treatment group, DBH slope values were consistently higher in the post-heatwave period. In the dry season, the DBH slope distribution during and post heatwaves showed a notably narrow interquartile range (IQR) in control trees, while the irrigated trees displayed a broader IQR.

### 3.3 Heatwave Impact

To assess how heatwave severity affects stem expansion and contraction, we employed a linear mixed-effects regression model. In this model, the DBH slope during heatwaves was modeled as a function of normalized heatwave severity, season (wet or dry), and treatment (control or irrigated), with individual trees included as a random effect. We focused on the wet and dry seasons, as they represent the two extremes of soil moisture availability, providing a clear contrast. The full model results are presented in Table S2. Marginal and conditional R^2^ values were 0.20 and 0.23, respectively.

For both seasons and treatments, higher normalized heatwave severity was associated with increased stem contraction rates, as indicated by negative DBH slopes of greater magnitude (Fig. 6a,b). The regression slope did not show statistically significant differences across seasons or treatments. In contrast, the intercept varied significantly: it was higher in the dry season for both treatments, with a much stronger increase observed for the irrigated plot. During the wet season (panel a), control and irrigated trees exhibited similar responses—mild stem contraction during weaker heatwaves, with contraction rates increasing as heatwave severity intensified. In the dry season (panel b), control trees followed a similar pattern, but irrigated trees displayed a qualitatively different response: their stems continued to expand during heatwaves, and greater heatwave severity only reduced the rate of expansion rather than reversing it into contraction.

**Figure 6:**
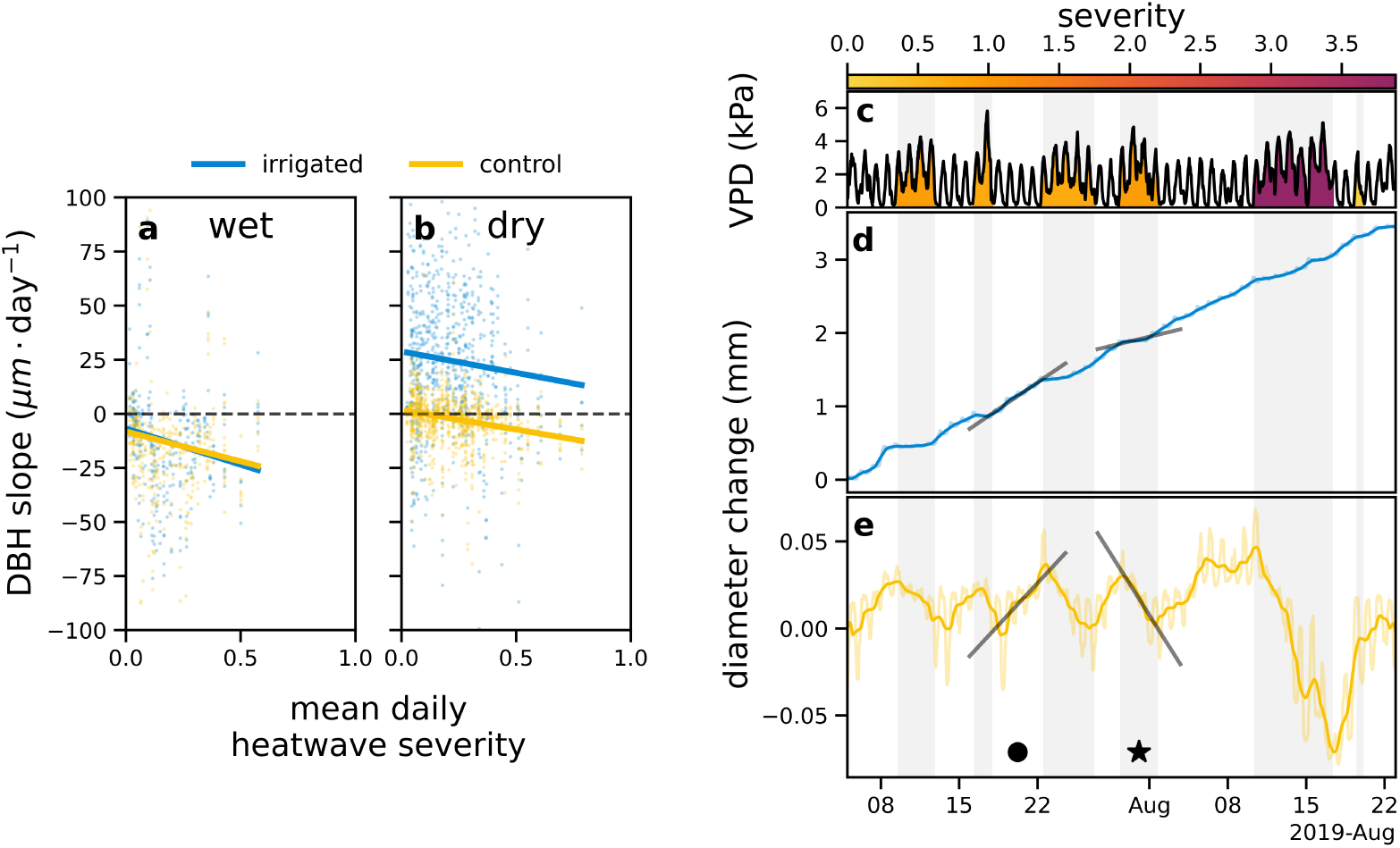
More severe heatwaves are associated with decreasing DBH slopes. Panels. **a,b:** Linear mixed model regression curves for wet and dry seasons and both treatments. Mean daily heatwave severity, treatment, and season were chosen as fixed effects, and individual trees were included as a random effect. Dots represent the response of individual trees to each heatwave. Coefficients for regression curves can be seen in Table S2. **Panels c–e:** Illustration of the impact of heatwaves on DBH slope, for two individual trees (one control and one irrigated). **Panel c:** VPD time series with shading indicating heatwave severity. Periods classified as heatwaves are also marked with a light gray background across panels c–e to provide contextual alignment. **Panels d,e:** DBH time series for an irrigated (d) and a control (e) tree. The light shaded colored lines denote half-hourly DBH data, while the thicker lines show the DBH trend, without the daily fluctuations. Black lines indicate the linear best fit for DBH during two distinct periods, one when a heatwave ocurred, marked with a star, and one post heatwave, marked with a circle. For the irrigated tree, the DBH slope was 104.2 µm *·* day*^−^*^1^ post-heatwave and 36.6 µm *·* day*^−^*^1^ during the heatwave. For the control tree, the DBH slope was 6.9 µm *·* day*^−^*^1^ post-heatwave and *−*10.2 µm *·* day*^−^*^1^ during the heatwave.

To illustrate this point, panels d and e show the stem diameter change of a representative irrigated tree and control tree over a 45-day period during the dry season. To provide context, panel c displays the corresponding VPD time series, with heatwave periods shaded according to their severity. When no heatwaves occur (e.g., the period marked with a black circle), both trees exhibit stem expansion, consistent with the pattern shown in Fig. 5n. In contrast, during heatwaves (e.g., the period marked with a black star), the irrigated tree continues to expand—albeit at a reduced rate—while the control tree exhibits stem contraction, in agreement with the results shown in Fig. 5m.

## 4 Discussion

### 4.1 Growth Rates

The comparison between the growth rates in the control and irrigated plots raises a few interesting patterns.

First, the harsh environmental conditions in the Yatir forest result in high stem growth rates during the wet (and colder) season. This contrasts with the typical seasonal growth pattern observed in pine trees (Maseyk, 2006; Maseyk et al., 2008; Rotenberg and Yakir, 2010), where peak growth generally occurs during the warm season. Interestingly, the irrigation experiment brings the pine trees closer to this usual growth pattern. Although phenology is not the main focus of this paper, it is important to note that not all tissues in the control trees grow best in the wet season. For instance, needles in control trees have been observed to reach their highest growth rates in June (Maseyk, 2006; Maseyk et al., 2008).

Second, we see that the growth of trees in the control plot seems to be primarily water-limited, while irrigated trees are not (Littell et al., 2008; Dudney et al., 2023). For the irrigated trees, water limitation became evident in 2021 when irrigation was reduced to 50% of PET, which, although sufficient to support growth during the drying and dry seasons, led to diminished Ω values and growth rates, shifting from the typical full growth to disrupted or even spurt growth modes. In contrast, control trees exhibit a clear water-limited response. During the drying and dry seasons, they shift into a stagnant-GRO mode, characterized by growth rates declining to near zero. During this period, T-REX initially rises (contraction mode), then stabilizes at a plateau (standstill mode) that persists as long as no precipitation occurs. This plateau coincides with the lowest xylem water potential values recorded throughout the year (Preisler et al., 2022). We hypothesize that during this phase, trees adopt an extremely conservative water-use strategy to avoid the risk of irreversible hydraulic failure. The stagnant-GRO mode implies minimal to no actual stem growth, though some limited growth may still occur even under hydraulic deficit conditions (Zweifel et al., 2016). Pine forests across a wide climatic range can exhibit plasticity in their growth patterns (Rotenberg and Yakir, 2010), adapting the timing of their peak carbon assimilation to the most favorable conditions. Findings from the irrigation experiment indicate that individual pine trees also have this plasticity, enabling fast adaptation to favorable conditions within weeks or months (Preisler et al., 2023). This is evidenced by the fast divergence between irrigated and control trees at the onset of the irrigation experiment, see Figs. 3 and 4.

Third, the analysis of the time series shows no evidence of irreversible damage to the control trees’ ability to grow following the prolonged dry season, during which they remain in standstill mode. As the wetting period begins, T-REX declines (recovery mode), and within 3–4 weeks, the trees transition into increasing-GRO mode, with growth rates rising to match those of the irrigated trees. This pattern is corroborated by measurements of net photosynthesis and transpiration reported by Preisler et al. (2023). If there were some impairment in plant hydraulics such as high percent loss of conductivity (PLC), we would expect sub-optimal growth rates compared to the irrigated trees, which serve as a standard for trees with minimized hydraulic damage. Wagner et al. (2022) showed that the hydraulic damage to the trees in the Yatir forest is mainly in their needles, and does not reach high embolism levels.

Fourth, we observed a clear alignment between the growth rates of the trees in the control plot and the flux tower NEP trends. Generally, when growth rates are positive, NEP suggests that the forest acts as a carbon sink, while near-zero growth rates indicate a shift toward the forest acting as a carbon source. This qualitative agreement reinforces our decision to use the rate of change in GRO as a proxy for actual tree growth in the Yatir forest.

The year of 2020 provided two interesting deviations from the patterns above, which we attribute to the above average precipitation of the 2020 hydrological year. This year registered 63% greater precipitation than the average of the 20 preceding years, putting it in the 95th percentile. The first deviation is that most trees in the control plot showed unusual positive growth rates during the dry period. We hypothesize that higher than average soil water availability could have sustained positive growth rates even during the dry season. The second deviation is that although irrigated trees were provided only with 80% of PET, their growth rates seem comparable to previous years. Here, two hypotheses come to mind: higher soil water availability could have offset the decreased irrigation amounts, or irrigation at 80% of PET simply does not trigger water limitation responses.

Lastly, we would like to raise our choice of dividing the year into seasons based on precipitation dynamics, instead of doing so based on measured soil water availability. In the spirit of this paper, we strived to extract maximal insight from easily-measurable variables. In this sense, precipitation rates are a simpler measure than soil moisture content, since the interpretation of the latter depends on site-specific factors such as soil surface, depth of measurement, soil texture and its spatial heterogeneity, soil stoniness, etc (Preisler et al., 2019). This choice is validated by the alignment between season classification and observed growth modes. In irrigated trees, full growth occurs during the dry season, and spurt growth during the wet. In control trees, we generally observe increasing-GRO modes during the wet season, contraction during the drying season, standstill during the dry season, and recovery during wetting. The time series of growth modes offers an effective and intuitive summary of the trees’ qualitative state. Importantly, this approach is independent of the absolute DBH length units and relies solely on time-dependent metrics (Ω and T-REX). We propose that such a framework will be especially useful for comparing tree states across individuals of different sizes, species, and environments.

### 4.2 Seasonal Responses to Atmospheric Events

The convergence of DBH slope distributions between control and irrigated trees during the wet season—across all atmospheric conditions—underscores the capacity of the two cohorts to exhibit similar growth dynamics when both soil and atmospheric conditions are similar (see Fig. 5, panels e–h). This alignment supports our earlier interpretation of comparable growth rates under high soil moisture availability and reinforces the idea of tree plasticity: when water is not limiting, even trees subjected to contrasting long-term treatments respond similarly. The consistent divergence in drier seasons, on the other hand, highlights the central role of soil water availability in shaping differences in growth responses.

Rainfall events consistently triggered stem expansion across all seasons and treatments, likely reflecting a combination of growth and rapid water uptake (see Fig. 5, panels c, g and k). The elevated DBH slopes during these events highlight the trees’ ability to grow and quickly restore internal water reserves. Some of the observed expansion may also result from bark swelling due to wetting, independent of actual growth (Oberhuber et al., 2020).

Outside the wet season, irrigated trees generally exhibited higher DBH slopes than control trees, reflecting the effect of increased soil water. This pattern held even during rain events, where the same rainfall had different outcomes. We hypothesize that in the irrigated plot, the water was readily available for uptake, while in the control plot, part of it may have served to replenish depleted soil storage potentially explaining the slightly lower slopes (see Fig. 5, panels c and k).

DBH slopes were consistently higher in the post-heatwave periods compared to the heatwave periods in the same season, reflecting relief from atmospheric stress. These post-heatwave conditions—characterized by below-median VPD— were generally more favorable for tree growth across both treatments. Importantly, high atmospheric demand during heatwaves imposes a toll on trees regardless of soil water status. However, the nature of this toll—whether it leads to reduced growth rates or a transition to contraction—depends on soil moisture availability and season. These findings raise important questions about the role of heatwave severity (not considered in the analysis presented in Fig. 5) in shaping DBH slope responses, particularly in driving the transition from expansion to contraction—an issue explored in more detail in the following subsection.

### 4.3 Heatwave impact

The regression analysis reveals a consistent pattern: as the mean daily heatwave severity becomes more intense, trees show increasingly reduced DBH slopes, indicating slower growth or faster contraction. This trend was similar across both treatments and seasons, suggesting that the effect of atmospheric stress on stem dynamics operates independently of soil water status. Specifically, increases in the mean daily severity of a heatwave were associated with proportional reductions in DBH slope, regardless of whether trees were in wet or dry conditions. While the model explained only a modest share of the overall variance (conditional R^2^ of 0.23)—pointing to the role of additional influencing factors—the negative relationship between heatwave severity and DBH slope was statistically robust (see Table S2.

The differences in intercepts from the regression model helps explain the contrasting DBH slope patterns observed during heatwaves in the wet and dry seasons (Fig. 5, panels e and m). In the wet season, both control and irrigated trees exhibited similar responses, with overlapping DBH slope distributions (Fig. 5e) and comparable sensitivity to increasing heatwave severity (Fig. 6a). In contrast, during the dry season, irrigated trees maintained positive DBH slopes—indicative of continued stem expansion—while control trees, that experienced compound drought, showed mild contraction (Fig. 6b). This divergence can be attributed to the elevated intercept observed for irrigated trees in the dry season, reflecting a higher baseline growth rate. We hypothesize that growth is the key factor driving this difference: in the dry season, irrigated trees continue to grow actively due to long photoperiod combined with high soil water availability, allowing them to absorb the impact of heatwaves through a reduction in growth rate rather than a full transition to contraction. As mean daily heatwave severity increases, this expansion slows but rarely reverses. In contrast, control trees experience little to no growth during the dry season and thus lack the buffer of ongoing expansion, making them more susceptible to contraction under compound drought. This highlights the critical role of baseline growth dynamics in mediating stem responses to extreme atmospheric conditions.

This pattern is clearly illustrated in the example shown in Fig. 6c–e. Over a 45-day period during the dry season, both a representative irrigated tree and a control tree exhibit stem expansion in the absence of heatwaves (black circle), reflecting background (post-heatwave) conditions. However, during heatwaves (black star), their responses diverge sharply: the irrigated tree maintains stem expansion, though at a slower rate (36.6 µm *·* day*^−^*^1^ vs. 104.2 µm *·* day*^−^*^1^), while the control tree shifts to contraction. This case highlights the buffering capacity of high baseline growth in irrigated trees, which allows them to remain in expansion mode even under stress. In contrast, the control tree, with minimal baseline growth (DBH slope of 6.9 µm *·* day*^−^*^1^), lacks this buffer and responds to atmospheric demand with net stem shrinkage. These individual trajectories reinforce the broader findings: soil water availability does not eliminate the negative effect of heatwaves, but it significantly alters the outcome—slowing growth instead of reversing it.

### 4.4 Last considerations

The analysis of DBH slope across seasons and atmospheric events (see Fig. 5 and Fig. 6) also included the later years of the irrigation experiment, 2020 and 2021, when irrigated trees received 80% and 50% of PET, respectively. Notably, the 2020 hydrological year also exhibited growth rate anomalies, likely related to unusually high precipitation (see Fig. 3). Despite these factors, the analysis still identified clear cases where trees in the control and irrigated plots differed significantly, highlighting the robustness of the observed patterns.

In this study, we focused on dry heatwaves, a common feature of Mediterranean climates. To characterize these heatwaves, we prioritized VPD over temperature, as VPD more directly reflects the atmospheric demand for water. Our analysis centered on the impact of VPD on tree water status and growth. However, extreme temperatures, which we did not investigate in this work, pose a significant threat to tree health and functioning. It is important to acknowledge that our findings do not necessarily apply in cases of extreme heat. High temperatures alone can cause irreversible damage to tree tissues, including mortality (Wahid et al., 2007; Teskey et al., 2015; Still et al., 2023).

## 5 Conclusion

In this study, we explored how pine trees respond to the pressures of soil and atmospheric droughts, with a particular focus on stem growth dynamics. By leveraging continuous dendrometer measurements from an irrigation experiment in a semi-arid Mediterranean forest, we disentangled the roles of soil water availability and atmospheric demand in shaping tree behavior across seasons and atmospheric events.

Our findings highlight the importance of soil water in sustaining growth and modulating responses to short-term atmospheric drought. When soil moisture was not limiting, trees—regardless of long-term treatment—exhibited similar growth dynamics, underscoring their physiological plasticity. In contrast, under soil drought, trees displayed divergent responses: irrigated trees maintained growth during heatwaves, while control trees experienced contraction.

To capture and interpret these dynamics, we utilized a suite of dendrometer-based metrics—GRO rate, Ω, and T-REX—that together provide a robust framework for assessing growth, contraction and recovery. Notably, Ω, and T-REX rely solely on stem diameter measurements and are unit-independent, making them broadly applicable across species, sizes, and measurement systems. GRO rate tracks irreversible stem expansion, Ω quantifies resistance through the fraction of time spent expanding, and T-REX captures recovery ability through the effective number of contraction days since irreversible growth last occurred. Together, these tools offer a multidimensional view of tree growth modes.

As climate change intensifies the intensity, duration and frequency of droughts, there is a growing need to better understand how trees respond to shifting water availability. This study offers a new lens for interpreting high resolution stem diameter variation data, revealing how growth dynamics—expansion, contraction, and recovery—unfold across a range of soil and atmospheric conditions. These insights enhance our ability to monitor tree functioning in real time, offering potential early warning indicators of stress and supporting efforts to track tree growth across natural, agricultural, and urban environments.

## 6. Acknowledgements

EF was supported by a PhD scholarship from the Israel Ministry of Environmental Protection and the Israel Council for Higher Education. DY and ER were supported by grants from the Israel Science Foundation (ISF), the Weizmann Institute SAERI/IES and, and the Keren Kaymet L’Israel (KKL-JNF). YM acknowledges the support of the Research Center for Agriculture, Environment and Natural Resources, of the Hebrew University of Jerusalem. We would like to thank Jonathan Friedman and Niv DeMalach for useful discussion.

## 7 Author contributions

CRediT author statement. **Erez Feuer:** Conceptualization, Methodology, Software, Formal analysis, Investigation, Writing - Original Draft, Visualization. **Yakir Preisler:** Conceptualization, Investigation, Writing - Review & Editing. **Eyal Rotenberg:** Writing - Review & Editing, Investigation, Resources. **Dan Yakir:** Writing - Review & Editing, Supervision. **Yair Mau:** Conceptualization, Methodology, Writing - Original Draft, Visualization, Supervision, Project administration.

## 8 Data availability

Data will be made available upon acceptance at Figshare (DOI link).

## Supporting Information

### VPD time series and heatwave severity

We show below the time series of VPD and VPD percentiles over a period of eight months.

**Figure S1:**
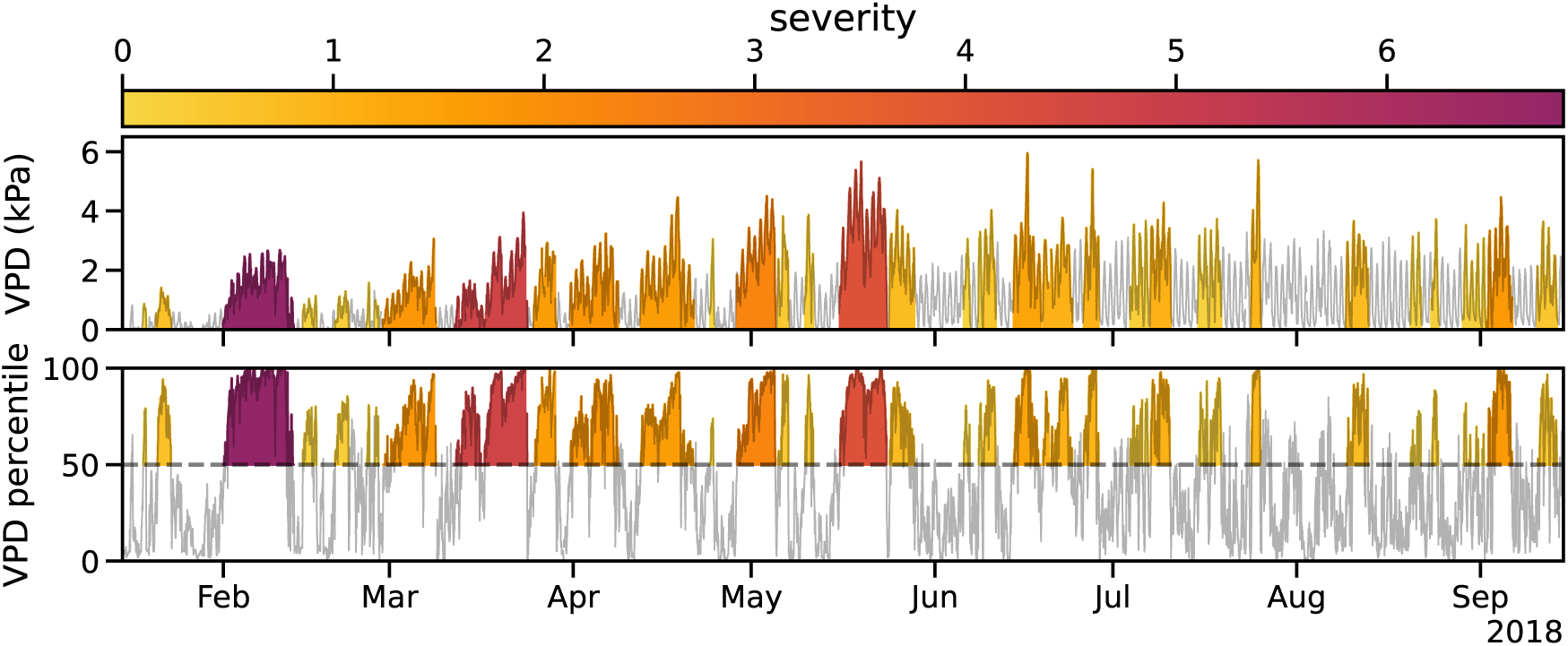
Heatwave classification and characterization. Heatwave periods are identified based on sustained VPD percentiles above the median. The severity of each heatwave (colors according to the colorbar on the top) is then quantified according to Eq. (1).

### T-REX algorithm

The algorithm below assumes zero-based indexing (first index of an array is zero), and in a “for loop” the last element is excluded. This is used in the Python language, but users of Matlab, Rlang, Fortran, etc, should be mindful of necessary adjustments.

#### Algorithm 1

Recursive T-REX Algorithm

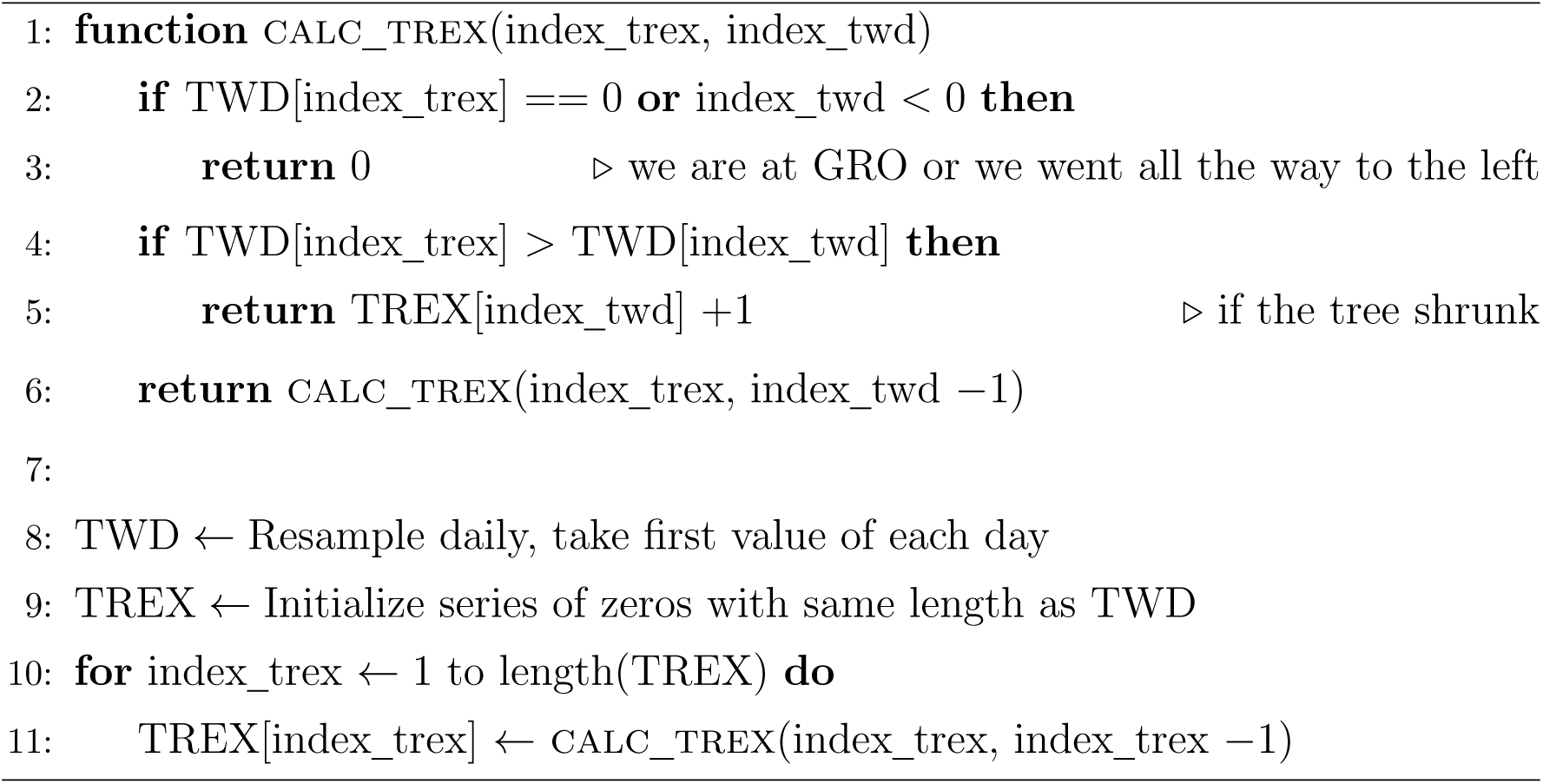

### Tables of the statistical analyses

**Table S1:** Statistical analysis of DBH slope responses across different events and seasons. This table presents the median differences (Δ) between trees in the control and irrigated plots during various atmospheric events (heatwave, post heatwave, rain, post rain) across all seasons (wetting, wet, drying, dry). Labels correspond to those presented in Figure 5. Adjusted p-values reflect multiple testing correction using the Benjamini–Hochberg procedure.

| Label | Season | Event | $\Delta$ | Adjusted p-value |
| --- | --- | --- | --- | --- |
| a | wetting | heatwave | 6.500 | 0.047 |
| b | wetting | post heatwave | -43.504 | 0.003 |
| c | wetting | rain | -35.540 | 0.033 |
| d | wetting | post rain | -10.250 | 0.011 |
| e | wet | heatwave | 1.046 | 0.752 |
| f | wet | post heatwave | -4.693 | 0.605 |
| g | wet | rain | 6.144 | 0.640 |
| h | wet | post rain | 3.058 | 0.243 |
| i | drying | heatwave | -16.699 | 0.006 |
| j | drying | post heatwave | -25.850 | 0.012 |
| k | drying | rain | -22.377 | 0.043 |
| l | drying | post rain | 0.877 | 0.738 |
| m | dry | heatwave | -25.329 | 0.003 |
| n | dry | post heatwave | -70.856 | 0.003 |

**Table S2:** Coefficients from the mixed-effects model analyzing heatwave impact on DBH slope. This table presents the estimated coefficients, p-values, and 95% confidence intervals (CI) for each parameter in the mixed-effects model. The model includes interactions between normalized severity, treatment (irrigated), and seasonal effects (wet vs. dry), along with the random effect for group variability (individual trees). The default season is wet and the default treatment is control, shown in bold. To obtain the coefficients of the non-default conditions as presented in Figure 6a,b, the coefficients should be added to the default, following this example: the coefficients for irrigated trees during the dry season are the sum of rows 1–4 for the intercept and rows 5–8, for the slope.

| Row | Parameter | Coefficient | p-value | CI Lower | CI Upper |
| --- | --- | --- | --- | --- | --- |
| 1 | **Intercept: wet, control** | -8.24 | 0.0227 | -15.33 | -1.149 |
| 2 | ΔIntercept: irrigated | 1.39 | 0.795 | -9.083 | 11.86 |
| 3 | ΔIntercept: dry | 10.07 | $8.25 \times 10^{-4}$ | 4.169 | 15.97 |
| 4 | ΔIntercept: irrigated & dry | 25.58 | $1.36 \times 10^{-8}$ | 16.75 | 34.41 |
| 5 | **Slope: wet, control** | -27.53 | 0.0132 | -49.31 | -5.751 |
| 6 | ΔSlope: irrigated | -5.82 | 0.728 | -38.58 | 26.94 |
| 7 | ΔSlope: dry | 9.28 | 0.468 | -15.79 | 34.36 |
| 8 | ΔSlope: irrigated & dry | -1.52 | 0.937 | -39.29 | 36.24 |
| 9 | Random Effect (Group Var) | 0.06 | 0.0344 | 0.004201 | 0.11 |

